# Bacterial diversity turnover estimates in a continental river system

**DOI:** 10.1101/2025.04.25.650558

**Authors:** Katalin Demeter, Domenico Savio, Alexander K.T. Kirschner, Georg H. Reischer, Stoimir Kolarevic, Juraj Parajka, Julia Derx, Stefan Jakwerth, Christian Wurzbacher, Alfred P. Blaschke, Robert L. Mach, Günter Blöschl, Andreas H. Farnleitner, Alexander Eiler

## Abstract

Understanding bacterial dynamics in large river systems is crucial for predicting continental-scale ecological functioning under anthropogenic pressures. Here, two consecutive surveys 6-years apart along the 2600 km Danube River found that carbon incorporation per cell and hour decreased by 5000 atoms every kilometer and that cells multiplied five times during their travel down the entire river. Resolving these cell turnovers taxonomically revealed taxa with a hundredfold difference from these average numbers. Bacterial community turnover was due to replacement (phylotype turnover) and could be linked to species sorting. This was despite an overall decrease in diversity richness downstream. Using linear models, we were able to relate carbon, cell, phylotype and diversity turnover rates to water residence time and discharge with outliers associated with human impacts. As such the reproduceable macroecological models predict microbial changes from anthropogenic and climate alterations along a continental drainage system providing insights into their ecological consequences.

## Main

Microorganisms play an integral and often unique role in the functioning of ecosystems and thus provide essential ecosystem services to human society. Gaining insight into the large-scale patterns of microorganisms – including their carbon, cell, species, and community turnover – is thus essential for predicting anthropogenic impacts on terrestrial and aquatic ecosystems. Macroecological studies on bacterial communities – including both experimental and large-scale surveys – have revealed that communities are assembled by ecological drift, dispersal-related processes, and local environmental conditions, in a process referred to as ‘species sorting’ ^1–4^. While these and other ^5–8^ studies have provided a theoretical framework to make rough predictions on the dynamics of the bacterial compartment, the question of the relationship between secondary production as well as cell, species, and community turnover along continental drainage networks is still insufficiently addressed, and actual rate estimates are largely missing.

In aquatic systems, the water residence time (WRT) is considered to be a main parameter in the community assembly process ^7–9^ and was shown to regulate carbon, cell, and biomass turnover ^10–12^. Particularly for aquatic environments with long WRT such as lakes, soils, alluvial groundwater aquifers, and the open ocean, the local environmental conditions have often been argued to be of uppermost importance in determining the composition of bacterial communities ^13–17^ and microbial functioning including bacterial secondary production ^10,18^.

With increasing river width and decreasing riparian influence, species sorting is considered to progressively prevail over allochthonous inputs of bacterial cells, *i*.*e*., ‘mass effects’, in shaping the bacterioplankton composition. For headwater streams, the generally high flow velocities and low WRT entail that the initial mass effects must exceed the rate of local extinction (i.e. export rate) to sustain cell densities as typically observed even in extremely pristine streams ^6,19^. In contrast, in large rivers with neglectable cell imports from tributaries, internal cell production must exceed the loss rates to explain the observed increasing cell densities ^20–23^. Cell production is thereby directly related to bacterial secondary production, which itself is limited by autochthonous carbon fixation as well as allochthonous input of carbon sources and nutrients ^10,12^. The latter include anthropogenic sources such as sewage, city runoff, or agricultural runoff. On a higher level, nutrient inputs, cell production, and losses, and therefore cell densities are governed by the prevailing hydrological conditions, and WRT in particular ^10,12,18^. Mass effects from tributaries and diffuse sources can vary locally and over temporal time scales ^24^.

Surface-runoff-induced riparian influence is mainly determined by precipitation events ^25^, while droughts represent conditions of minimum riparian influence. Consequently, hydrological and meteorological conditions in the catchment regulate both the internal production and the mass effects; and thus determine bacterial biomass as well as organic matter transformation and concentrations ^26–29^. While temporary fluctuations in the hydrological regime in response to meteorological events have been shown to affect the dynamics and observable distribution of bacterial taxa ^30,31^, these observations have so far not been put into the context of macroecological dynamics. Quantitative models may allow us to predict microbial dynamics, such as secondary production, and cell, genotype, and community turnover, occurring in the hydrological path from the continents to the ocean in light of climate change and other anthropogenic pressures.

For modeling purposes, the effects of the hydrological regime (i.e. WRT and discharge) on bacterial species and community turnover need to be comprehended in the context of bacterial production (i.e. cell division rates or doubling times) and the standing stock (i.e. cell abundances and biomass), and thus cell turnover rates. Typical bacterial concentrations in streams and rivers are reported to range from 10^5^-10^8^ cells per mL ^20,32^ with daily cell division rates ranging from 0.1 to 2 ^10,21,33,34^. By calculating the doubling times (calculated as 1/division rate) under a given WRT, we can estimate the number of generations on which species sorting can act in a system ^10^. Combined cell, species, and community turnover rates are essential for the modeling of temporal and spatial variations in microbiological water quality and nutrient budgets over continental scales. This is of particular importance considering that rivers represent the major link from the continents to the ocean in the global water cycle, as well as many ecosystem services such as drinking water by river bank filtration, sources for irrigation water, transport routes for shipping, and contaminant degradation.

Here, we hypothesize that changes in individual bacterial phylotypes as well as in community composition depend on water residence times and bacterial generation times which in turn are highly influenced by climate and anthropogenic modification to the continental drainage systems. As a study site, we selected the 2600 km-long Danube River (**Fig. S1**), the second largest river in Europe by discharge and length which drains a basin of approximately 801 000 km^2^ belonging to 19 countries ^35^. We compare the datasets of two whole-river surveys supplemented by data from a monthly sampling campaign over one year at two sites. This unique setting allows studying the upstream-to-downstream devolvement of bacterial community composition over a flow time (water residence time) of several weeks. We hypothesized that this long travel time, coupled with the warm water temperatures during the surveys, allows for sufficient bacterial cell divisions to observe species sorting. We estimated the number of bacterial cell divisions and the total cell production during the travel time based on measurements of bacterial secondary production and cell counts. Additionally, we analyzed bacterial diversity based on 16S rRNA gene sequences. Observations on the effect of discharge and anthropogenic factors, such as impoundments and wastewater treatment plants, complement this comprehensive study.

We report turnover rates from two large consecutive surveys (*Joint Danube Surveys*; short *‘JDS’*) that were performed in a highly similar fashion along the ~ 2600-kilometre-long shippable main branch of the Danube River (**Fig S1**) six years apart (JDS2 in 2007 and JDS3 in 2013). The results show reproducible macroecological patterns and provide a basis for the modeling of large-scale, short-as well as long-term changes in response to changing hydrological conditions. These conditions can be floods, droughts, and water scarcity, all predicted to increase under ongoing climate change.

## Results and Discussion

### A heavily impacted, continental river

The river discharge indicated mean flow conditions during JDS2 in 2007 and prevailing low flow conditions during JDS3 in 2013 (**Fig S3**, ^36^). The course of the Danube River is heavily modified by river regulation and damming along its entire course, with the two Iron Gate Reservoirs (RO, SRB, **Fig S1)** being the most significant impoundments. This results in highly varying flow velocities along the hydrological path (ranging from 0.6 to 11.2 km h^−1^ during JDS2 and from 0.9 to 5.4 km h^−1^ during JDS3, **Fig S3**). Since the surveys were conducted during summer, water temperatures were on average 21.1 °C during JDS2 and 20.9 °C during JDS3. Nitrate concentrations showed an initially synchronously decreasing trend (from 13.8 to 3.9 mg L-^1^ in JDS2 and from 12.0 to 3.9 mg L-^1^ in JDS3), while a slight increase was observed after the Iron Gates for JDS2 (up to 7.3 mg L-^1^). Orthophosphate concentrations did not show any clear longitudinal trends, though having lower concentrations during JDS2 (mean 0.11 mg L-^1^) than JDS3 (mean 0.15 mg L^−1^) six years later. The abundance of phytoplankton, as indicated by chlorophyll-a measurements, showed stark regional differences, indicating point sources of nutrient input between the two major cities Budapest and Belgrade (ranging from 0.7 to 30.6 µg L-^1^during JDS2 and from 0.3 to 16.0 µg L-^1^ during JDS3, **Fig S3**, ^36^).

### Trends in bacterial cell concentrations

Total bacterial cell concentrations – or “Total Cell Counts” (*TCC*), enumerated using epifluorescence microscopy (Experimental Procedures) – showed a clear overall increasing trend along the river (**Tables 1 and 2**; **Fig. 1A**). The ~ 10-fold discrepancy in cell concentrations between the two surveys is likely not only due to different hydrological conditions but also the result of a change in methodology in counting cells across the two surveys (Experimental Procedures). Independently from the methodological bias, the upstream-to-downstream increase in cell concentrations aligns well with another previous investigation along the Danube River ^21^ as well as with reports from other river systems such as the Australian Murray River ^22^ and the river Rhine ^23^.

**Table 1:** Summary table of descriptive statistics on measured and calculated parameters for both surveys.

| Parameter | Unit | Data type | JDS2<br>Median(range) | JDS3<br>Median(range) | Other studies |  |
| --- | --- | --- | --- | --- | --- | --- |
| cumulative travel time, $tt_{cum}$ | days | calculated | 33.7 | 49.7 | | |
| $TCC$ | $10^9$ cells $L^{-1}$ | measured | 1.71 (0.72 – 5.10) | 13.9 (2.98 – 26.2) | | |
| $BSP$ | $\mu gC L^{-1} h^{-1}$ | measured | 0.32 (0.14 – 1.46) | 1.15 (0.07 – 3.57) | 0.95 – 4.83<br>1.97 – 3.23<br>0.7 – 81 | Danube River <sup>21</sup><br>Danube River <sup>33</sup><br>Mississippi River <sup>37</sup> |
| $CD_d$ | $day^{-1}$ | calculated | 0.12 (0.03 – 0.92) | 0.12 (0.009 – 0.47) | 0.09 – 1.04 | Danube River <sup>21</sup> |
| $CD$ | cell divisions / ~ 2600 km | calculated | 4.2 | 6.0 | | |
| $aCP_d$ | $10^6$ cells $L^{-1} km^{-1}$ | calculated | 3.02 (0.77 – 18.0) | 35.1 (4.6 – 145) | | |
| $CP_{tot}$ | $10^9$ cells $L^{-1}$ | calculated | 9.21 | 95.9 | | |

TCC: total cell count, BSP: bacterial secondary production, CD: cell division rate, aCP: absolute cell production.

**Table 2:** Summary table of linear models relating bacterial parameters to travel distance or travel time for both surveys. The probability of an error by rejecting the null hypothesis of no change over distance/time is < 0.05 in all models listed, except for the ‘Simpson diversity (between consecutive sites)’ model for JDS3 where it is >0.05.

| Parameter | Unit | JDS2<br>slope ± std. error | JDS2<br>n | JDS2<br>adj. $R^2$ | JDS3<br>slope ± std. error | JDS3<br>n | JDS3<br>adj. $R^2$ |
| --- | --- | --- | --- | --- | --- | --- | --- |
| $TCC$ change | cells $L^{-1} km^{-1}$ | $0.54 \pm 0.14 \times 10^6$ | 75 | 0.165 | $4.71 \pm 0.76 \times 10^6$ | 54 | 0.411 |
| $BSP$ change | $ngC L^{-1} h^{-1} km^{-1}$ | $-0.19 \pm 0.05$ | 75 | 0.185 | $0.34 \pm 0.11$ | 54 | 0.133 |
| $BSP_c$ change | $10^{-21} g C cell^{-1} h^{-1} km^{-1}$ | $-139 \pm 30$ | 75 | 0.215 | $-56 \pm 10$ | 54 | 0.383 |
| ACE richness (alpha diversity) | ASV $h^{-1}$ | $-0.145 \pm 0.046$ (FL)<br>$-0.176 \pm 0.070$ (PA) | 39<br>47 | 0.197<br>0.107 | $-0.112 \pm 0.018$ | 54 | 0.411 |
| Simpson diversity (between consecutive sites) | $10^{-4} h^{-1}$ | $-1.14 \pm 0.55$ (FL)<br>$-1.86 \pm 0.66$ (PA) | 40<br>52 | 0.081<br>0.123 | $-2.31 \pm 0.26$ | 53 | -0.004 |
| Simpson diversity (among all sites) | $10^{-4} h^{-1}$ | $4.84 \pm 0.21$ (FL)<br>$3.17 \pm 0.21$ (PA) | 820<br>1378 | 0.397<br>0.153 | $4.59 \pm 0.07$ | 1431 | 0.737 |

FL: free-living fraction (0.2–3.0 $\mu m$ ) of JDS2 samples, PA: particle-associated (>3.0 $\mu m$ ) fraction of JDS2 samples

**Figure 1.**
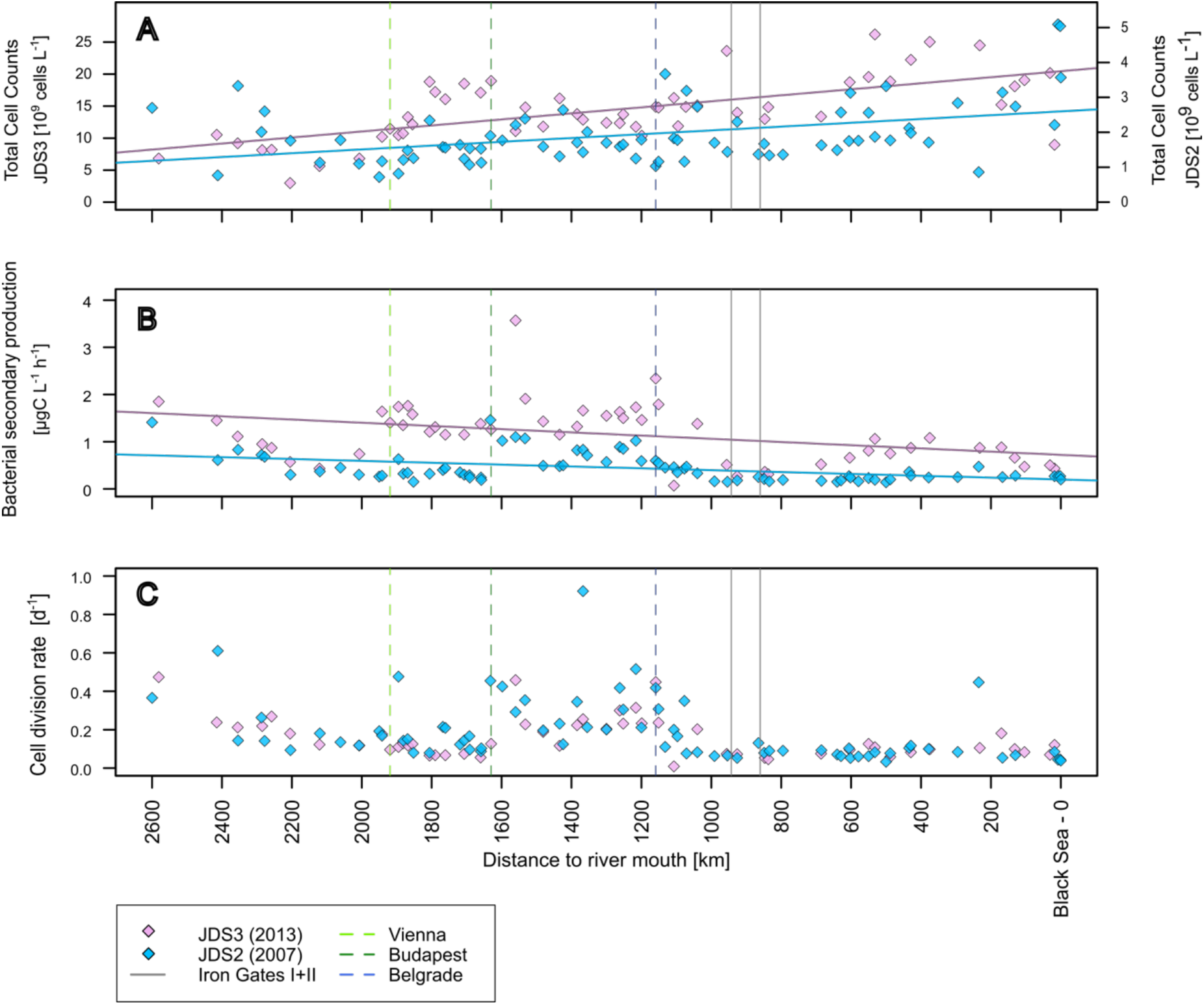
Trends in prokaryotic cell concentration (Total Cell Counts; TCC) (A), bulk bacterial secondary production (BSP) (B), and daily cell division rates (CD_d_) (C) along the Danube River from upstream (left; rkm 2600) to the river mouth (right; rkm 0). Regression statistics are shown in Table 2. (Data for JDS2 was taken from ^20^). n(JDS2) = 75, n(JDS3) = 54

### Trends in bacterial secondary production

Bacterial secondary production (*BSP*) is the rate of biomass production of a heterotrophic population measured by the incorporation rate of a carbon compound (Experimental Procedures). BSP of the bulk prokaryotic community along the river ranged from 0.14 to 1.46 μgC L^−1^ h^−1^ with a mean of 0.44 μgC L^−1^ h^−1^ during JDS2 (^20^, **Table 1, Fig. 1B**), and from 0.07 to 3.57 μgC L^−1^ h^−1^ with a mean of 1.16 μgC L^−1^ h^−1^ during JDS3 (**Table 1, Fig. 1B**). These *BSP* rates – serving as the basis for subsequent calculations of doubling times and cell division rates (*CD*_*d*_) – were in a typical range as previously reported for the Danube River, but also other river ecosystems (**Table 1**). General linear models to estimate overall trends along the river showed a significant decrease in *BSP* by 0.19 ± 0.05 ngC L^−1^ h^−1^ km^−1^ (JDS2) and 0.34 ± 0.11 ngC L^−1^ h^−1^ km^−1^ (JDS3), (**Table 2**), although pronounced local increases were observed in river sections with strong anthropogenic impact.

During both surveys, elevated *BSP* values were observed in the section of the three large capital cities with > 1 million inhabitants, starting around rkm 1895 at Vienna (AT) over Budapest (HU) till rkm 1040 downstream of Belgrade (SR) (**Fig. 1B**). The higher BSP values were probably facilitated by phytoplankton growth due to sewage (nutrient) inputs from the large cities (as indicated by chlorophyll-a concentrations, ^20^ and **Fig. S3**). This correlation between *BSP* and phytoplankton growth has also been observed during the first Joint Danube Survey conducted in 2001 ^38^. The pronounced decrease in *BSP* downstream of Belgrade in the backwater of the first Iron Gate reservoir (rkm 1040) was accompanied by decreasing chlorophyll-a and TSS concentrations and increasing zooplankton abundance (Rotifers, Cladocera & Copepods; ^39^). This increase in grazing and particle sedimentation is likely a consequence of large-scale damming at the Iron Gates. A second peak in *BSP* was observed during JDS3 in the lower reach of the river (rkm 686-132). This, however, could not be explained by phytoplankton growth but is likely directly related to cell and nutrient import from highly polluted tributaries with generally very high *BSP* rates ^20^.

Cell-specific bacterial secondary production (*BSP* per cell; *BSP*_c_) showed a similar pattern as observed for bulk bacterial production with a modeled overall decrease of 139 ± 30 (adj. R^2^= 0.21, p < 0.001) and 56 ± 10 ? 10^−21^g C cell^−1^ h^−1^ km^−1^ (adj. R^2^= 0.38, p < 0.001) over the entire sampled longitudinal river transect during JDS2 and JDS3, respectively (**Fig. S4A**). This corresponds to a modeled decrease of 6900 and 2800 carbon atoms incorporated per cell per hour every kilometer, for JDS2 and JDS3, respectively.

### Enough time for species sorting

The average cell doubling time (i.e. time for doubling of a single cell, **Fig S2A**, Formula Collection) was estimated for each station based on bulk *BSP* rates and mean cell biomasses calculated from cell volumes by fluorescence microscopy (m*BM*_c_; ranging from 15 to 55 × 10^−15^ gC cell^−1^ for JDS2 and 6 to 36 × 10^−15^ gC cell^−1^ for JDS3). Median cell doubling times were 8.05 (JDS2) and 8.23 (JDS3) days, corresponding to median daily cell division rates (*CD*_*d*_) of 0.124 (JDS2) and 0.121 (JDS3) divisions per day (**Fig. 1C, Table 1**). The obtained values were within the range of values reported from the Danube River more than 25 years ago^21^ (**Table 1**). Based on the calculated median *CD*_*d*_ for the bulk communities, this allows for 4.2 and 6.5 cell divisions (i.e. doubling events) for a cell on its estimated ~ 33.7 and ~ 53.6-day long journey along the investigated ~ 2600 km longitudinal river section for JDS2 and JDS3, respectively. Actual *CD*_*d*_ values certainly vary among bacterial taxa and the presented values are best estimates.

In the context of WRT, the calculated doubling times are important quantitative measures for the understanding of ecological processes in aquatic systems. The average in-stream WRT, i.e., the time it takes for an average parcel of water to move through the river, thereby is the most applicable predictor when discussing the development of native as well as allochthonous bacterial populations (i.e., soil and groundwater) being exposed to varying riverine conditions. The WRTs or travel times per kilometer themselves underly variation both along the course of a river and over time (see the section below), being influenced by discharge dynamics as well as by anthropogenic impoundments, which in turn affect ecological processes.

### Total cell production along the river

The absolute cell production per river kilometer and liter (a*CP*_*km*_) was estimated between consecutive sampling sites based on the mean bulk *BSP* between these sites (m*BSP*_up⟷down_), the mean biomass of cells at the upstream site (m*BM*_c up_) and the flow distance between these sites (*d*_*up*⟷*down*_, **Fig S2B**, Formula Collection). a*CP*_*km*_ ranged from 0.77 × 10^6^ to 18.0 × 10^6^ cells L^−1^ km^−1^ during JDS2 and from 4.61 × 10^6^ to 144.9 × 10^6^ cells L^−1^ km^−1^ during JDS3, and were on average higher by a factor of 9.9 during JDS3 than during JDS2 (**Fig S4B**). By summing up these numbers for each section between two sites, the total cell production in one liter of river water traveling along the entire river (*CP*_*tot*_) was estimated to be 9.2 × 10^9^ (JDS2) and 95.9 × 10^9^ (JDS3) cells. Putting these numbers in relation to the median standing stocks along the river as determined by microscopic quantification (median TCC, 1.7 × 10^9^ cells L^−1^ for JDS2 and 13.9 × 10^9^ cells L^−1^ for JDS3, **Fig 1A**), the total cell production along the river equals ~ 5.4 (JDS2) and ~ 6.9 (JDS3) times the median standing stock, meaning that the cell loss rates compensating for this production and therefore necessary for the observed cell counts (*TCC*) must be in the same range. Since our calculations did not account for allochthonous inputs such as tributaries and anthropogenic point sources (WWTPs, sewer overflows, direct sewage discharge in lower reaches), out of which especially the latter is probably a net source of cell input, real cell loss rates along the river must be even higher.

Taxa with cell division rates below the average daily cell division rates (*CD*_*d*_) of 0.124 (JDS2) and 0.121 (JDS3, see section *‘Enough time for species sorting’*) will not be able to compensate for the average cell losses. These taxa will therefore not be able to sustain their population along the river, and would therefore decrease in absolute abundance. Yet, this is only true under the simplified assumption that loss rates are constant and equal among individual taxa and species. Preferential predation by bacteriophages and heterotrophic nanoflagellates or removal by filter feeders due to specific cell size characteristics will for example affect species-specific cell losses, and thus bacterial species sorting and composition ^40,41^.

The above discussed cell replacement (cell production and consequently cell loss) rates - i.e., that the median cell stock is replaced at least 5.4 (JDS2) and 6.9 (JDS3) times during the travel time along the river - theoretically allows for, but not necessarily entails high bacterial community turnover rates (high beta diversity).

### Diversity and community turnover

16S rRNA gene amplicon sequencing data from the two surveys conducted six years apart showed highly corresponding trends in bacterial alpha (species/phylotype richness within a sample) and beta diversity (differences in species/phylotype composition among samples) along the entire river, indicating universal, continuous (hydrological) processes that are responsible for shaping the bacterial communities (**Fig. 2A and B**). Given the importance of the available time for the division of bacterial cells and thus for species sorting, estimates of alpha and beta diversity changes along the river are modeled as a function of travel time in the river.

**Figure 2.**
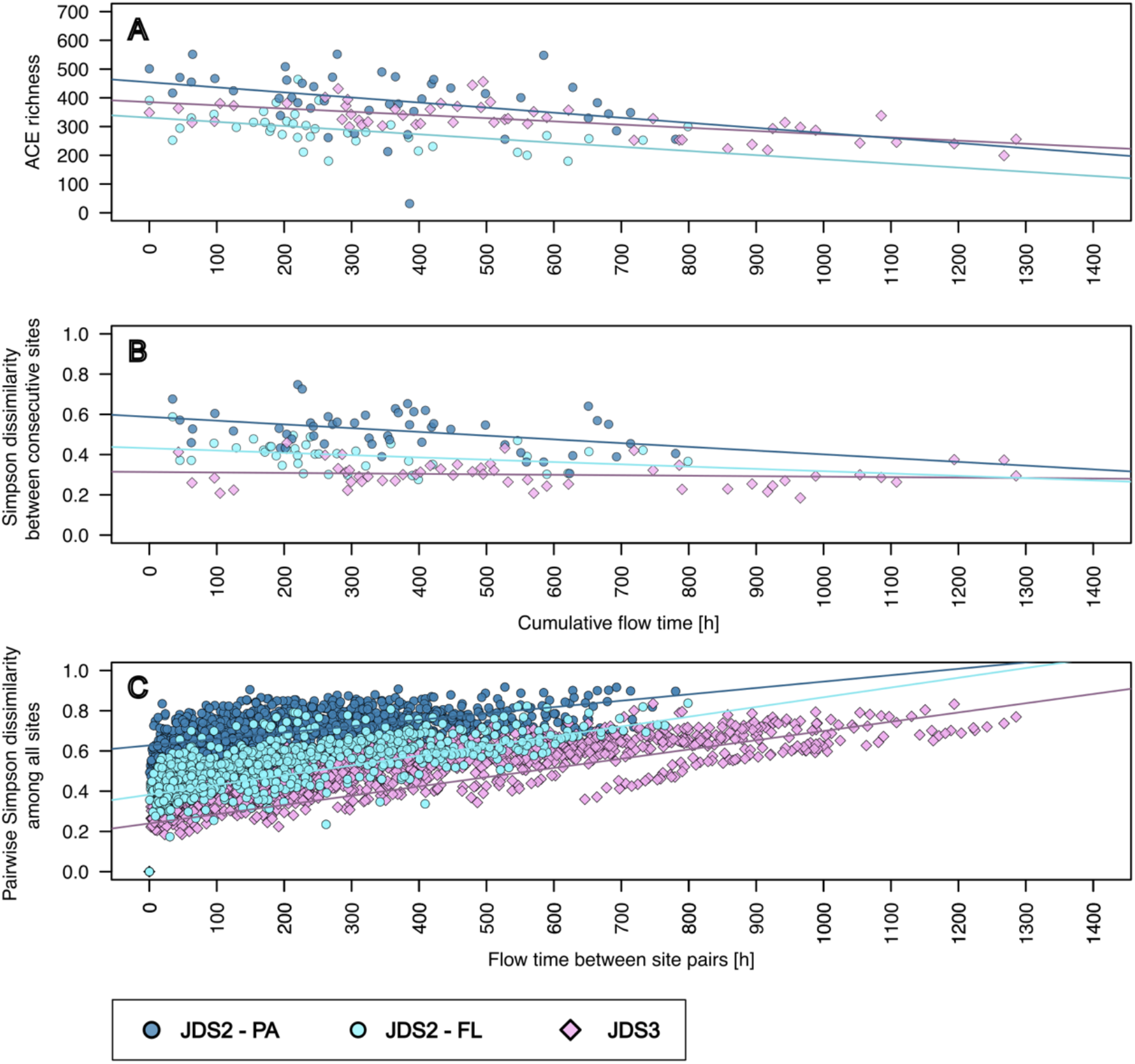
The gradual development of alpha and beta diversity with increasing cumulative travel time (*tt*_*cum*_) during JDS3 (total community; violet; n = 54) and for the two separated size fractions studied during JDS2, representing the bacterioplankton communities of 0.2-3.0 µm (light blue; n = 39) and >3.0 µm (dark blue, n = 47) - corresponding to free-living and particle-associated bacterioplankton, respectively. Panel **(A)** depicts the development of alpha diversity (ACE richness), **(B)** the dynamics of phylotype turnover (phylotype replacement) component of beta diversity between two consecutive sites given as Simpson dissimilarity, and **(C)** the pairwise phylotype turnover component among all sites. The colored lines represent fitted linear models for the respective datasets. Regression statistics are shown in Table 1. JDS2-FL: free-living fraction, JDS2; JDS2-PA: particle-associated fraction, JDS2.

Similarly to the pioneering study on bacterial diversity along the Danube River during JDS2 ^7^, JDS3 also showed a gradual decrease in alpha diversity (ACE richness) along the river (**Fig. 2A, Table 2**). This coincided with a gradual development of the bacterial communities as shown for all three datasets (JDS3; JDS2 free-living fraction; JDS2 particle-associated fraction) in the NMDS visualization of beta diversity analysis results (**Fig. S5)**. The model for alpha diversity revealed a decrease (phylotype loss) of 0.14 ± 0.05 amplicon sequence variants per hour [ASVs h^−1^] in the free-living community (JDS2-FL) and 0.18 ± 0.07 ASVs h^−1^ in the particle-associated community during JDS2 (JDS2-PA). During JDS3, the alpha diversity decreased by 0.11 ± 0.02 ASVs h^−1^ (**Fig. 2A, Table 2**).

Beta-diversity, as determined by Simpson dissimilarity, was modeled to be 4.84 ± 0.21 × 10^−4^ h^−1^ (JDS2-FL), 3.17 ± 0.21 × 10^−4^ h^−1^ (JDS2-PA) and 4.59 ± 0.07 × 10^−4^ h^−1^ (JDS3) (**Fig. 2B, Table 2**), thus changing by a factor of 0.426 ± 0.017 (JDS2-FL), 0.250 ± 0.015 (JDS2-PA) and 0.592 ± 0.01 (JDS3) over the entire sampled continental river transect. Variations in community turnover (represented by the projections of bacterial community samples in the Bray-Curtis based non-metric multidimensional scaling (NMDS, **Fig. S5**) were found to be statistically significantly correlated to geomorphological measures such as ‘distance to river mouth’ (river kilometer), ‘mean dendritic stream length’ (as a proxy for in-stream WRT) and catchment area, with remaining variations related to total nitrate, chlorophyll a, total suspended solids and phosphorus (**Table S1**).

Beta-diversity, i.e., community turnover, can be partitioned into two components, *nestedness* and *turnover*, based on the basic mechanisms driving differences in community heterogeneity among sites: the loss of phylotypes and their replacement, respectively ^42,43^. Estimates of phylotype (ASV) turnover given as rate varied between 0.18 and 0.75 between successive sites along the river continuum, with an overall trend of lower turnover towards the river mouth (**Fig. 2B**). Corresponding nestedness ranged from 0.001 to 0.397 with a median of 0.045 (free-living JDS2), 0.055 (particle-associated JDS2) and 0.036 (JDS3) between consecutive sites along the river continuum. Overall, phylotype turnover along the entire river was estimated to be between 0.92 and 0.96, contributing a by far larger part to overall beta diversity (ranging from 0.94 to 0.97 as estimated by Sorensen dissimilarity) when compared to nestedness which varied between 0.009 to 0.017. As ASV turnover exceeds nestedness, beta-diversity patterns in the Danube River mainly resulted from phylotype replacement processes rather than absolute phylotype loss. Estimates close to one further emphasize an almost complete exchange of phylotypes throughout the continental drainage system under low base flow conditions, coinciding with the previous observation of an increase of freshwater- and lake-bacteria ^7^.

### Re-occurring growth patterns

As a verification of the models for bulk secondary production *(BSP)* and thus cell turnover rates (sections ‘*Cell division rates and travel time frame species sorting’* and ‘*Cell replacement rates allow for high bacterial community turnover rates’*), we assessed individual phylotype changes. To achieve this, we estimated absolute phylotype abundances using relative abundances as retrieved from amplicon sequencing along with the *TCC* for each sample. Using these estimates, we observed ASVs with an increase in abundance by a factor of more than 100, corresponding to 6.6 cell division events or generations (the average number of cell divisions along the Danube River was 4.2 and 6.0 in JDS2 and JDS3, respectively). ASVs affiliated with the phylum *Actinobacteria* showed particularly large increases in relative abundance, growing from zero or nearly zero up to 16% of the total community along the river (ASV_3 during JDS3), reaching an absolute abundance of 2.3*10^8^ cells L^−1^ (**Fig. 3**). **Table S2** gives an overview of minimum and maximum relative and absolute phylotype abundances. **Fig. 3** also shows the fitted generalized additive models that estimate the realistic range of *in-situ* growth and loss rates of specific ASVs that naturally occur along the investigated large river. Interestingly, the calculated cell division rates (*CD*_*d*_) necessary to explain the observed dynamics are significantly lower (10 times) than the growth rates reported from a very recent study isolating 20 different bacterial strains from a polluted river in Belgium ^44^. This discrepancy might be due to environmental factors or biases of culture-based approaches selecting mainly for bacteria with high growth rates and neglecting natural loss factors such as predation by grazing and viral lysis.

**Figure 3.**
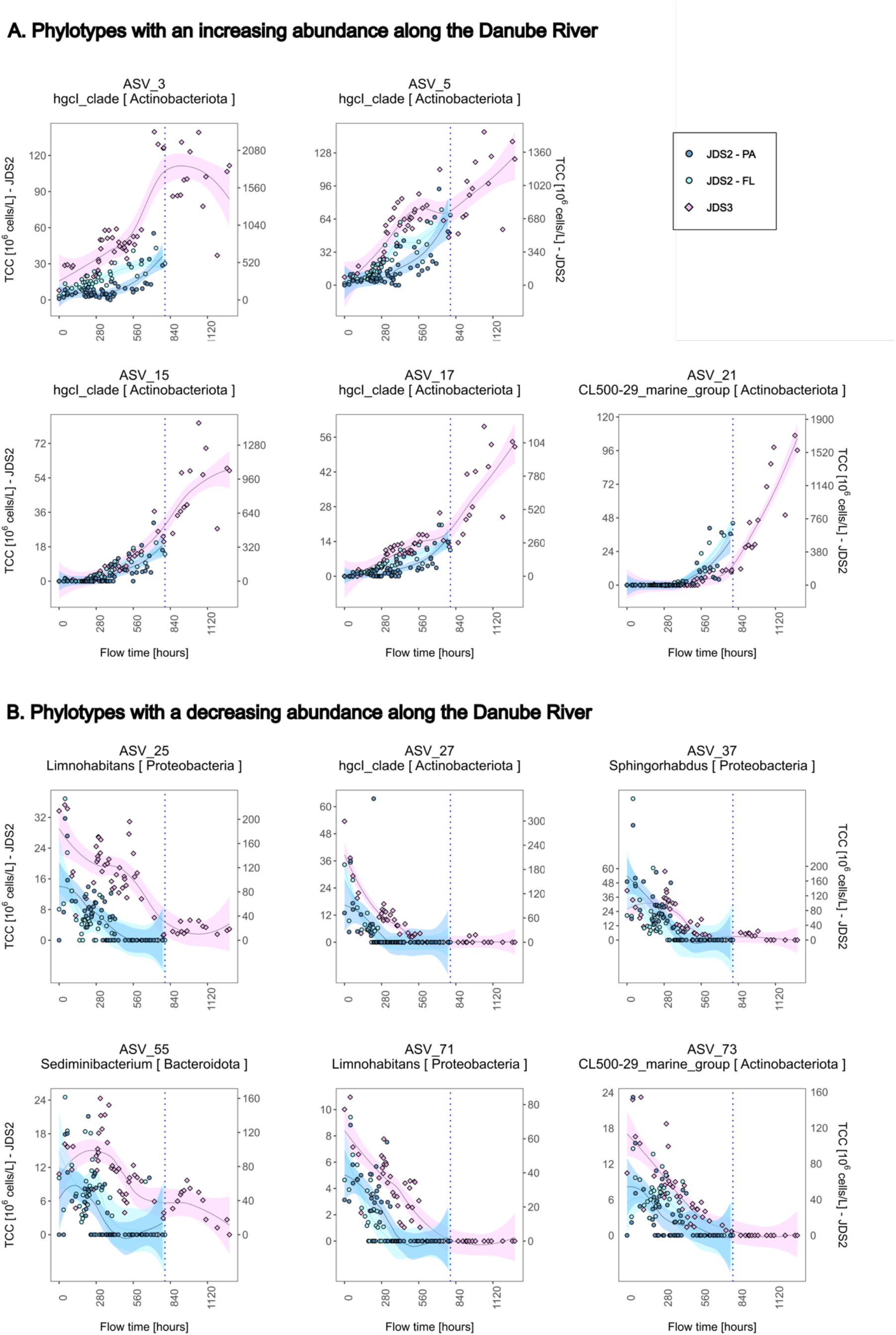
Changes in the abundance of selected phylotypes during their travel time along the Danube River. The lines show the fitted generalized additive models (R package *mgcv*), the ribbon marks their Bayesian credible interval (similar to confidence interval). JDS2-FL: free-living fraction, JDS2; JDS2-PA: particle-associated fraction, JDS2.

In contrast to these highly competitive phylotypes, we also observed phylotypes with significantly decreasing abundances along the river. These included - amongst others - ASV_25 and ASV_71 affiliated with genus *Limnohabitans*, ASV_37 of the genus *Sphingorhabdus*, ASV_73 of the ‘CL500-29 marine group’, ASV_27 affiliated with the hgcI clade in the phylum Actinobacteria and ASV_55 o the genus *Sediminibacterium* (**Fig. 3**). Their highest relative abundance, at the upper stretches of the Danube, reached 4.4.% (ASV_27 during JDS3), corresponding to an absolute abundance of 3*10^8^ cells L^−1^, and decreased to zero by the middle or the lower stretches. These decreasing trends can either be a result of species sorting due to lower competitiveness, or of higher specific loss rates (grazing, sedimentation, predation by phages). Besides positive selection due to lower taxon-specific grazing pressure (e.g. of small cells not filtered as efficiently by filter feeders), favorable environmental conditions with regard to nutrient status or cell import from tributaries may play major roles as indicated by an association between community and environmental dynamics along the river (**Table S1**). For example, the rise of ASV_15, ASV_17 and ASV_21 affiliated to the phylum *Actinobacteria* coincided (ASV_15) or followed (ASV_17 & ASV_21) alterations in environmental conditions along the Danube River due to damming at the iron gate hydropower plants (**Fig. 3**). These barriers create the largest reservoir along the Danube with considerable effect on flow dynamics by creating an impoundment of around 300 km upstream ^36^.

### Discontinuities in longitudinal trends

Aside from the modeled general trends along the longitudinal river transect, we also observed local discontinuities in particular for bacterial numbers *(TCC)*, mean cell-specific biomass (m*BM*_*c*_), bacterial secondary production *(BSP)* as well as for phylotype and community turnover rates. For example, *BSP* was highest in the section between the three large cities, Vienna (rkm 1919), Budapest (rkm 1660-1630) and downstream Belgrade (rkm 1159; **Fig. 1B**). The highest *CD*_*d*_ *rates*, however, were observed between Budapest (rkm 1630) and Belgrade (rkm 1159; **Fig. 1C**). Elevated *TCC* and *BSP* downstream of Vienna (rkm 1919), Budapest (rkm 1,630), Belgrade and in the section between rkm 629 and rkm 500 along the Romanian-Bulgarian border can be speculated to be associated with intensive agriculture and wastewater input from municipalities and/or agronomy. Major changes in the dynamics of individual species and community turnover along the river could be observed in association with the major cities, the Iron Gates, or confluences with tributaries.

In addition to the two longitudinal surveys, a monthly monitoring was conducted at both large cities (Vienna & Belgrade) over the period of one year. This yearly cycle revealed a positive response of alpha diversity (ACE richness) to increasing discharge. Normalizing the discharges measured at each sampling date to the minimum discharge observed at the respective site (‘discharge ratio’) allowed a comparison of the two sites with different discharge regimes. The generalized linear models of discharge-dependent response in ACE richness for the two monitoring sites predict that a doubling in discharge results in an increase in richness of 111 ± 46 ASVs at Vienna and 148 ± 33 ASVs at Belgrade (**Fig. 4**). Whether this estimate derived from a linear model can be generalized to the entire Danube River or even other continental drainage systems has to be subject to future studies, which also have to investigate additional factors explaining the large residuals. High-resolution data of the hydrological history (i.e. whether sampling was at the beginning or rather the end of a flood event), likely affecting the extent of cell mobilization from the riparian zone ^30,31,45^, could explain parts of the residuals in alpha diversity. Nonetheless, our result is supported by previous observations that locally increasing discharge levels as a consequence of snowmelt or precipitation events in a sub-catchment cause increasing species richness due to higher groundwater input and surface runoff ^9,31,45^.

**Figure 4.**
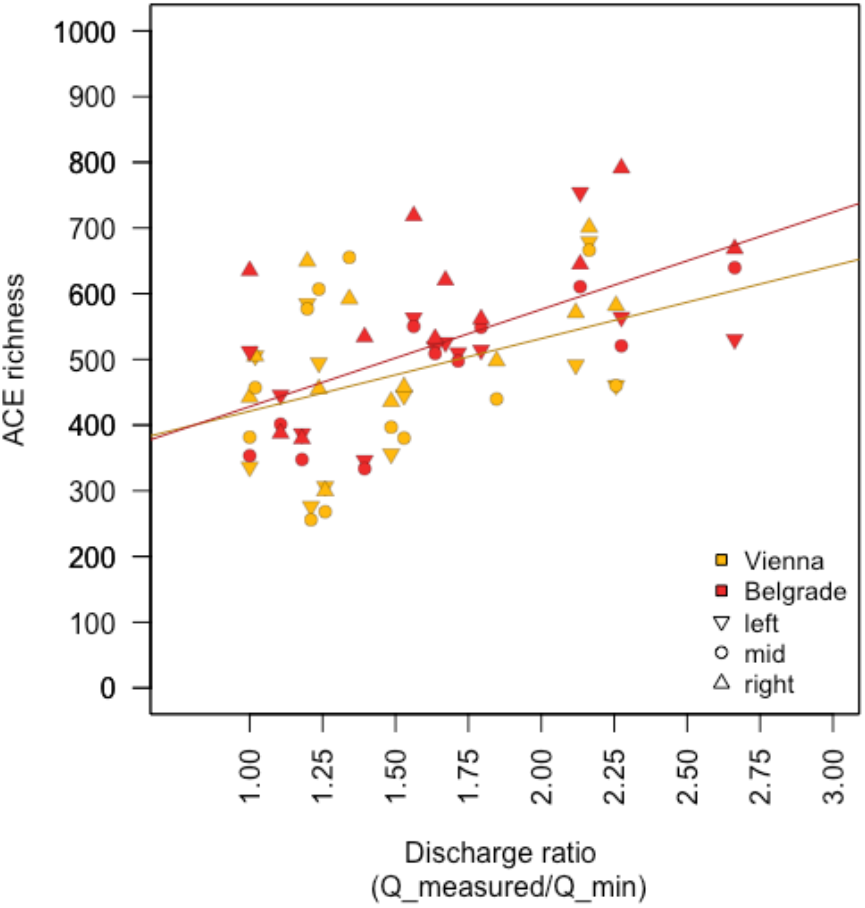
Correlation between alpha diversity (ACE richness) and prevailing discharge levels during the 15-month monitoring of bacterial community dynamics at two large cities along the Danube River from January 2014 - January 2015 in Vienna [AT], rkm 1919 and Belgrade [RS], rkm 1159. The x-axis depicts the ratio between the prevailing discharge at the respective sampling date normalized to the minimum discharge measured at the respective site during the entire monitoring. Regression statistics are shown in Table 1.

## Conclusions

The essential role of the hydrogeological structure in lotic aquatic systems - representing the link between land and the ocean in the global water cycle - in controlling macroecological processes has been suggested and debated in numerous studies. Here, we report highly reproducible patterns in bacterial secondary production as well as cell, phylotype, and community turnover rates along the large Danube River, allowing us to predict and model macroecological patterns of these bacterial turnover rates along a continental drainage system. The linear models allowed us to make some general predictions such as increasing cell numbers, decreasing secondary production and decreasing alpha diversity (phylotype richness) with increasing flow time. Apart from these overall trends, we observed local discontinuities in total cell counts and bacterial secondary production, possibly associated with locally intensive agriculture and wastewater input from municipalities. We also observed an increase in phylotype richness as a result of increasing discharge at two selected sites, during the annual cycle.

Beta diversity, i.e., community turnover was consistently associated with the replacement of phylotypes, rather than their loss. The stark increases and decreases in individual phylotypes, underlying phylotype replacement, were demonstrated through a selection of ASVs where the increasing ones not only reached the expected ‘growth’ from the calculated average cell division rates and hence generation times but also exceeded these.

We show that the main community-shaping mechanism along the river could simply be the number of possible cell divisions as a result of in-stream WRT. As such, climate warming resulting in decreased flow velocities associated with less streamflow in eastern Europe ^46^ or the construction of dams will translate into a quantifiable decrease in alpha diversity and enhanced community change along the river. Hence the bacterial diversity reaching the river’s estuary and the Black Sea will decrease, as previously proposed for another river system ^47^. Our findings highlight the power of hydrological measures such as discharge and WRT (travel time) to predict bacterioplankton (meta)community dynamics along the hydrological path from land to sea.

## Acknowledgments

This study was supported by the Austrian Science Fund (FWF) as part of the DKplus “Vienna Doctoral Program on Water Resource Systems” (W1219-N22) and the FWF projects P25817-B22 and P32464-B, as well as the research project “Groundwater Resource Systems Vienna” in cooperation with Vienna Water (MA31). AE was funded by the Swedish Foundation for Strategic Research (ICA10-0015) and the Swedish Research Council (VR2012-4592). Infrastructure (cruise ships, floating laboratory) and logistics for collecting, storing, and transporting samples were provided by the International Commission for the Protection of the Danube River (ICPDR). The analyses were performed using resources provided by Uninett Sigma2 AS under project “nn9744k”. We thank Bettina Premm for her contribution to the laboratory analyses. We also thank viadonau - Österreichische Wasserstraßen-Gesellschaft mbH - and the Federal Ministry of Agriculture, Regions and Tourism (BMLRT) of the Republic of Austria for the provision of discharge data for the Austrian river section for the years 2014 and 2015 via https://ehyd.gv.at.

## Materials and Methods

### Experimental Procedures

#### A comprehensive river survey for continental drainage systems

Within this study, we analyzed data from two large river surveys along the continental Danube River – namely *Joint Danube Survey* (*JDS*) 2 & 3 – conducted in the summers of 2007 and 2013. The overall purpose of the Joint Danube Surveys is to produce a comprehensive evaluation of the chemical and ecological status of the entire Danube River on the basis of the European Union Water Framework Directive (WFD), as well as the assessment of the microbiological status of the river ^48^. Within the scope of JDS3 (2013), in total, more than 800 individual chemical, microbiological, ecotoxicological, radiological and biological parameters were investigated ^36,49–51^. During JDS2 (2007), more than 280 individual parameters were determined ^7,20,48^. Data on bacterial secondary production, community dynamics and environmental parameters were published previously ^7^ and re-used in this study for in-depth calculations and comparison between JDS2 and JDS3.

#### Data and code availability

All data, sampling methods as well as analytical methods of Joint Danube Surveys 1, 2 and 3 are publicly available via the official website of the International Commission for the Protection of the Danube River (ICPDR; http://www.icpdr.org/wq-db/), and the final scientific report in particular ^36^. Selected data from JDS 1, 2 and 3 were published previously in several studies ^36,48,49,52–55^. All data analyses were performed in R. The R script and the corresponding workspace containing all data used for this manuscript is available at [link to github or Code Ocean].

#### Experimental procedures during Joint Danube Survey 2

The study design and experimental procedures of Joint Danube Survey 2 for results that are re-analysed in the current study were published by Savio, et al. ^7^.

#### Study sites and sample collection during Joint Danube Survey 3

Within the frame of JDS3 conducted between 13 August and 25 September 2013, water samples were collected in the midstream at 54 sampling stations along the longitudinal profile of the Danube River ^36^ (**Fig. S1**). This covers the shippable way from river kilometer (rkm) 2581 at Ulm/Böfinger Halde (Germany) to rkm 18 close to the river’s mouth at the Black Sea (Romania). Aside from the longitudinal snap-shot study, two sites (downstream of the cities of Vienna and Belgrade) were sampled monthly over approx. one year in 2014.

For DNA analysis, duplicate samples (biological replicates) were collected from midstream in sterile 1 L glass flasks from a water depth of approximately 30 cm. Glass flasks were sterilized by rinsing with 0.5% HNO_3_ and autoclaving. Depending on the expected biomass concentration, 120-300 mL of river water were filtered through 0.2 µm pore-size polycarbonate filters (Cyclopore, Whatman, Germany) by vacuum filtration for biomass concentration. The filters were stored at −80 °C until DNA extraction.

#### Geomorphological measures and estimation of in-stream water residence time

**Geomorphological parameters** including ‘catchment area’, ‘mean dendritic stream length’ and ‘cumulative dendritic distance upstream’ were calculated for each sampling site as described previously ^7^. The ‘mean dendritic stream length’ at a respective sampling site gives the average flow distance travelled by a drop of water through the entire catchment area to a particular sampling site (under the assumption that spring discharges are randomly distributed in the catchment), and therefore serves as a proxy for the average residence time in the stream (assuming constant flow velocities). In contrast, the ‘cumulative dendritic distance upstream’ was calculated as the sum of all mapped flow paths to a respective sampling site and represents a geometric parameter indicative of the drainage density, but not of residence times ^7^.

In contrast, **in-stream water residence time (‘travel time’)** along the main stream of the Danube River (*i*.*e*., without tributaries) from the very upstream site (Ulm, Germany) to the river mouth (Black Sea, Romania) was also estimated by an alternative approach based on on-site measurements of mean flow velocities at each sampling site. Here, **‘cumulative travel times’** for JDS2 and JDS3 were calculated by cumulating the calculated flow times between each two consecutive sampling sites. These, in turn, were estimated on the basis of averaged on-site 3D flow velocity measurements from two consecutive sampling sites and the flow distance between them. These calculations resulted in estimated ‘cumulative travel times’ of ~ 33.7 days and ~ 49.7 days from rkm 2600 to rkm 18 and rkm 2581 to rkm 18 for JDS2 and JDS3, respectively.

#### Bacterial numbers, biometry and bacterial secondary production

**Total prokaryotic cell concentration (*TCC*)** as well as **mean cell volumes (m*V***_***c***_) were determined by fluorescence microscopy as described previously by ^20^, with the difference that for JDS3, cells were stained using Sybr Gold fluorescence dye (Invitrogen SYBR™ Gold, Thermo Fisher Scientific; MA, USA) and counted on a ‘Nikon Eclipse 80i’ epifluorescence microscope, while for JDS2, Acridine orange (Merck, Darmstadt, Germany) staining dye and a ‘Leica Diaplan’ epifluorescence microscope were used. Moreover, for JDS2, cells were categorized as coccus-, rod- and vibrio-shaped cells for both the free-living and particle-associated filter fractions ^20^, while during JDS3, no separation of the two size fractions was made and cells were categorized as either ‘small’ or ‘large cells’ regarding counting and morphometry. ‘Small cells’ were defined as all coccus-shaped cells with a cell diameter of < 0.33 µm. All coccus-shaped cells with larger diameter as well as all cells not being coccus-shaped were categorized as ‘large cells’.

**Bacterial secondary production (*BSP*)** was determined for the bulk microbial communities based on ^3^H-leucine incorporation as described previously ^20^. For more detail on calculations see the ‘Formula collection’ and **Fig. S2**.

#### DNA extraction & 16S rRNA gene amplicon library preparation

Genomic DNA from JDS3 samples was extracted as for JDS2 using a slightly modified protocol of a previously published phenol-chloroform and bead beating-based procedure ^56^ using isopropanol instead of polyethylene glycol for DNA precipitation. Total DNA concentration was assessed applying the Quant-iT™ PicoGreen® dsDNA Assay Kit (Life Technologies Corporation, USA). 16S rRNA gene concentrations in the DNA extracts were quantified using domain-specific quantitative PCR as described previously ^7^. DNA extracts were normalized with regard to 16S rRNA gene concentrations in order to use standardized numbers of bacterial 16S rRNA gene templates for amplification and barcoding in a two-step barcoding procedure. In short, the first-step primers (341f and 805r) contained adapters for introducing Illumina adapters, and dual barcodes were used in the second step. The first-step PCR primers were thus (adapter sequence, followed by primer sequence) adapter-341f (‘5-ACACTCTTTCCCTACACGACGCTCTTCCGATCTNNNNCCTACGGGNGGCWGCAG-3’) and adapter-805r (‘5-AGACGTGTGCTCTTCCGATCTGACTACHVGGGTATCTAATCC-3’). The first-step amplicon PCR (ampPCR) was carried out in duplicate in 20 µL reaction mixtures containing 1 × Q5 reaction buffer, 0.2 mM dinucleoside triphosphates (dNTPs), 0.5 µmol L^−1^ forward and reverse primers, 0.4 U of Q5 high-fidelity DNA polymerase (New England BioLabs), as well as environmental DNA as template that was normalized to equal amounts of 16S rRNA gene copies prior to barcoding in order to increase comparability and reduce PCR bias. For normalization purposes, 16S rRNA gene copy concentrations in DNA extracts were determined by quantitative PCR as previously described ^7^. Cycling conditions for 1st-step ampPCR were 98 °C for 1 min, followed by 20 cycles of 98 °C for 10 s, 62 °C for 30 s, 72 °C for 30 s, and a final extension at 72 °C for 2 min. The duplicate products were pooled and purified using the Agencourt AMPure XP purification system (Beckman Coulter). The second PCR step containing variable combinations of primers with different multiplex-identifiers for sample-specific ‘barcoding’ (forward, AATGATACGGCGACCACCGAGATCTACAC-[index]-ACACTCTTTCCCTACACGACG; reverse, CAAGCAGAAGACGGCATACGAGAT-[index]-GTGACTGGAGTTCAGACGTGTGCTCTTCCGATCT) binding to the first-step adapters and incorporating Illumina adapters was carried out in single 20 µL reactions. Reactions contained 1 × Q5 reaction buffer, 0.2 mmol L^−1^ dinucleoside triphosphates (dNTPs), 0.25 µ mol L^−1^ forward and reverse index primers, 0.4 U of Q5 high-fidelity DNA polymerase (New England BioLabs) and 2 µL of purified amplicons from 1st-step ampPCR as template. Cycling conditions for the 2nd-step index PCR (idxPCR) were 98 °C for 1 min, followed by 15 cycles of 98 °C for 10 s, 66 °C for 30 s, 72 °C for 30 s, and a final extension at 72 °C for 2 min. PCR products (amplicon libraries) were purified as described above and quantified with the PicoGreen kit (Life Technologies). Products were sequenced at the SciLifeLab SNP/SEQ sequencing facility at Uppsala University, Uppsala, Sweden, on an Illumina MiSeq (2 × 300 bp) in two runs. A subset of the JDS2 samples (n=66) were processed twice, as technical replicates. Technical replicates showed high similarity (results not shown, analyses available in the supplementary R code).

#### 16S rRNA gene amplicon data analysis

Previously published (JDS2; ^7^) as well as newly generated raw sequence data (JDS3) were processed separately. For JDS3, raw amplicon sequencing data from two sequencing runs were automatically demultiplexed by the Illumina MiSeq sequencing software before primers were removed using the CUTADAPT tool ^57^ and sequences without matching primers were discarded. For the re-analysis of JDS2 samples, previously published and deposited amplicon libraries for 150 samples were downloaded from NCBI Sequence Read Archive (accession number SRP045083, ^7^). The R package dada2 (^58^, version 1.8) was used for de-replication, denoising, and sequence pair concatenation. After manual inspection of quality score plots, the forward and reverse reads of the bacterial 16S rRNA gene amplicons were trimmed at 260 and 200 bp length, respectively. Additional quality filtering removed any sequences with unassigned base pairs and reads with a single phred score below 10. After de-replication of reads, forward and reverse error models were created in dada2 with a subset of the sequences (~ 10^7^ sequence reads). Chimeras were removed using the ‘removeBiomeraDenova’-function provided in the R package ‘dada2’. Taxonomy was assigned using the Bayesian classifier and SILVA non-redundant database 138 ^59,60^. Next, the biological replicate pairs for the JDS3 dataset were merged and averaged to obtain the final amplicon sequence variant (ASV) table (**Datasets S1 and S2**). Chloroplast, cyanobacterial, mitochondrial, eukaryotic as well as archaeal ASVs were removed resulting in 10,801 (JDS2) and 12,417 (JDS3) ASVs. The biological replicates showed high similarity (results not shown, analyses available in the supplementary R code).

For the analysis of trends in **beta diversity** - the variation in species composition among sites - based on Bray-Curtis dissimilarity index, the sequence read abundances were first normalized to 1 for each sample using the ‘drarefy’-function implemented in the R-package ‘vegan’ ^61^. The Simpson dissimilarity is bounded between 0 and 1, where 0 means the two sites have the same composition (that is they share all the species and the species have the same abundances), and 1 means the two sites do not share any species. For the partitioning of beta diversity in regard to the respective turnover (i.e. species replacement between two sites) and nestedness (i.e. species loss between two sites) components, functions implemented in the R package ‘betapart’ were used ^43^.

**Alpha diversity** was described by the ACE-richness estimator. Prior to ACE-richness calculation, libraries with < 3,155 sequence reads were discarded, while the remainder were randomly resampled with replacement in 50 consecutive repetitions using the ‘rrarefy’-function implemented in the R package ‘vegan’. After averaging of the results from 50 repetitions and rounding, the mean number of reads of sequence reads was 3,152 reads per sample.

#### Statistical analyses

Statistical analyses and plot generation were conducted in R version 3.6.2 ^62^. Linear regression models were calculated using the ‘lm’ function implemented in R, reporting an ‘adjusted R squared’ as measure for the ‘goodness of fit’ and the Standard Error as a measure for variation. ASV abundance changes were modelled using the R packages ‘mgcv’ ^63^ and ‘gratia’ ^64^. Generalized additive models (gam) with simple factor smoothers on rkm were fitted to all ASVs (in terms of estimated cell concentrations) being detected in at least three samples. The fitted curves were then differentiated, using the derivatives function implemented in the ‘gratia’ package ^64^, to estimate the net changes of each ASV. From the fitted derivative curves we calculated maximum absolute either decrease or increase for each ASV.

### Formula Collection

This section complements Figure S2.

#### A. Doubling time and cell division rate (for each site)

##### Bacterial numbers and biometry

Mean cell volumes per cell (m*V*_*c*_) [µm^3^ cell^−1^] were determined by averaging calculated cell volumes from at least 100 randomly selected cells per sample as determined by epifluorescence microscope-based morphometry by measuring the diameter (cocci) or length and widths (all other morphotypes) for each cell and calculating volumes either as spheres (for cocci), cylinders with two half spheres at the ends (rods, curved rods, filaments). Cell volumes for single cells were calculated as follows:

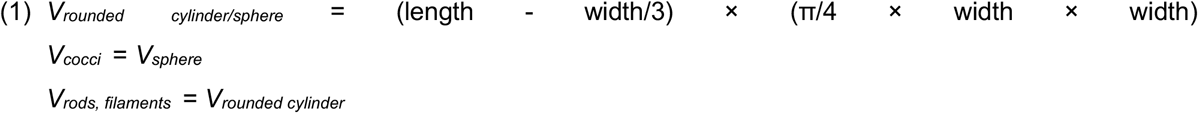

**Mean cell volumes** (m*V*_*c*_) [µm^3^ cell^−1^] for each site was calculated as the weigthed mean considering each cell type (cocci, rods, curved rods and filaments in JDS2; small and large cells in JDS3).

**Mean cell biomass** (m*BM*_*c*_) [fgC cell^−1^] was computed for each site by multiplication of calculated mean cell volume at the respective site (m*V*_*c*_; in µm^3^) with a commonly applied dry weight conversion formula of 120 × m*V*_*c*_^0.72^ - representing the functional allometric relationship between cell biomass (in femtograms C) and mean cell volume (in cubic micrometers; m*V*_*c*_) of bacteria ^65^.

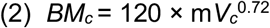

##### Bacterial secondary production

**Bacterial secondary production** (*BSP*) [µgC L^−1^ h^−1^] of the bulk prokaryotic communities was determined based on ^3^H-Leucine incorporation rates (LI) [mol L^−1^ h^−1^],

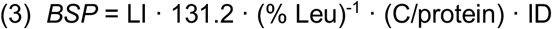

where the constant of 131.2 represents the molar mass [g mol^−1^] of leucine, % Leu is the fraction of leucine in protein (0.073) ^66^, C/protein is the ratio of cellular carbon to protein (0.86), and ID is the isotope dilution. For ID, the value of 1 was used based on leucine uptake kinetics experiments conducted in parallel with sample measurements. These were conducted on representative samples from along the river in order to determine the concentration at which the uptake of ^3^H-Leucine was saturated, and no isotope dilution was observed ^20^. Based on these experiments, a final concentration of ^3^H-Leucin of 100 nmol L^−1^ was selected for both surveys.

**Bacterial secondary production per cell** (*BSP*_c_) [fgC h^−1^ cell^−1^] was calculated for each sampling site by dividing the total *BSP* [µgC L^−1^ h^−1^] by the total cells counts (*TCC*) [cells mL^−1^] as determined with epifluorescence microscopy.

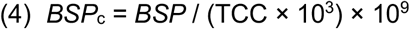

##### Doubling time and cell division rate

**Doubling times** (*DT*_*d*_,) [days] were calculated by dividing the mean cell biomass (m*BM*_*c*_) [fgC cell^−1^] by the mean

*BSP* per cell (*BSP*_*c*_) [fgC h^−1^ cell^−1^] at the respective site.

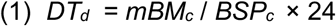

**Daily cell division rates** (*CD*_*d*_) [day^−1^] for each sampling station were calculated as the reciprocal of daily doubling times (*DT*_*d*_).

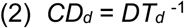

#### B. Cell production rates (per travel distance or travel time)

**Absolute cell production rates per hour** (a*CP*_*h*_) between two consecutive sites [cells mL^−1^ h^−1^] were calculated by dividing the mean *BSP* of the bulk bacterial communities of two sites (m*BSP*_up⟷down_) [µgC L^−1^ h^−1^] by the mean biomass per cell at the upstream site (m*BM*_*c up*_) [fgC cell^−1^]. To obtain cell production rates per mL and hour, m*BSP*_up⟷down_ had first to be multiplied by 10^9^ and then be divided by 10^3^ to obtain production in fg C per mL:

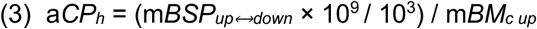

**NOTE**: In case no *BSP* measurements were available for neither site, the median *BSP* from all sites was used.

**Absolute cell production between two consecutive sites** (a*CP*_*up*⟷*down*_) [cells mL^−1^] was estimated by multiplication of a*CP*_*h*_ with the calculated travel time in hours (*tt*_*up*⟷*down*_) [h] between two consecutive sites.

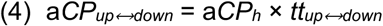

**Travel time between two consecutive sites** (*tt*_*up*⟷*down*_) in hours [h] was estimated by dividing the flow distance between two sites (*d*_*up*⟷*down*_) in km by the mean flow velocity (m*v*_*up*⟷*down*_) [m s^−1^] between the two sites as estimated by averaging the measured flow velocities at the two sites.

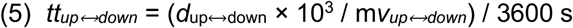

Cumulative travel times in days [d] for the two surveys were estimated by simply summing up all between site travel times:

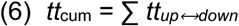

**Absolute cell production rate per km** (a*CP*_*km*_) between two consecutive sites [cells mL^−1^ km^−1^] was calculated by dividing the calculated absolute cell production between each two sites (a*CP*_*up*⟷*down*_) by the flow distance between two sites (*d*_*up*⟷*down*_).

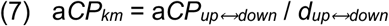

Estimated **total cell production along the entire river** (*CP*_*tot*_) [cells mL^−1^] during both surveys was calculated by summing up all calculated cell production values between all sites.

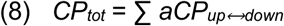

## Notes

**Conflict of Interest Statement:** The authors declare no conflict of interest.

### Competing Interest Statement

The authors have declared no competing interest.

