## Supplementary Figures and Tables for "Bacterial diversity turnover estimates in a continental river system"

**This document includes:**

Figures S1 to S5
Tables S1 to S2
SI References

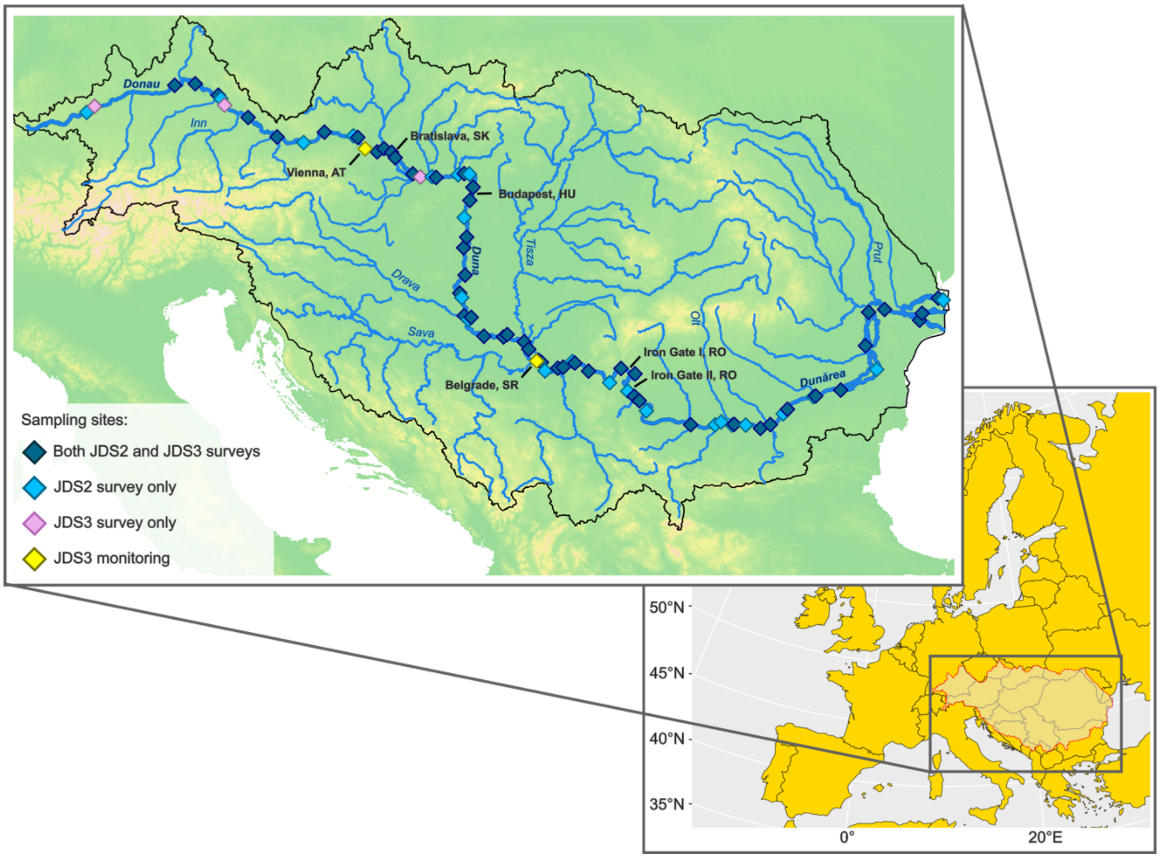


**Figure S1.** Map of the Danube River catchment illustrating the sampling sites during the Joint Danube Survey 2 (JDS2, taken from (Savio et al., 2015), n = 75), the Joint Danube Survey 3 (JDS3, n = 54) and the one year monthly monitoring campaign following JDS3 (n = 2). Country capitals along the Danube and the two largest dams along the river, namely Iron Gate I and II, are depicted in black font. The map in the background shows the Danube River catchment within Europe.

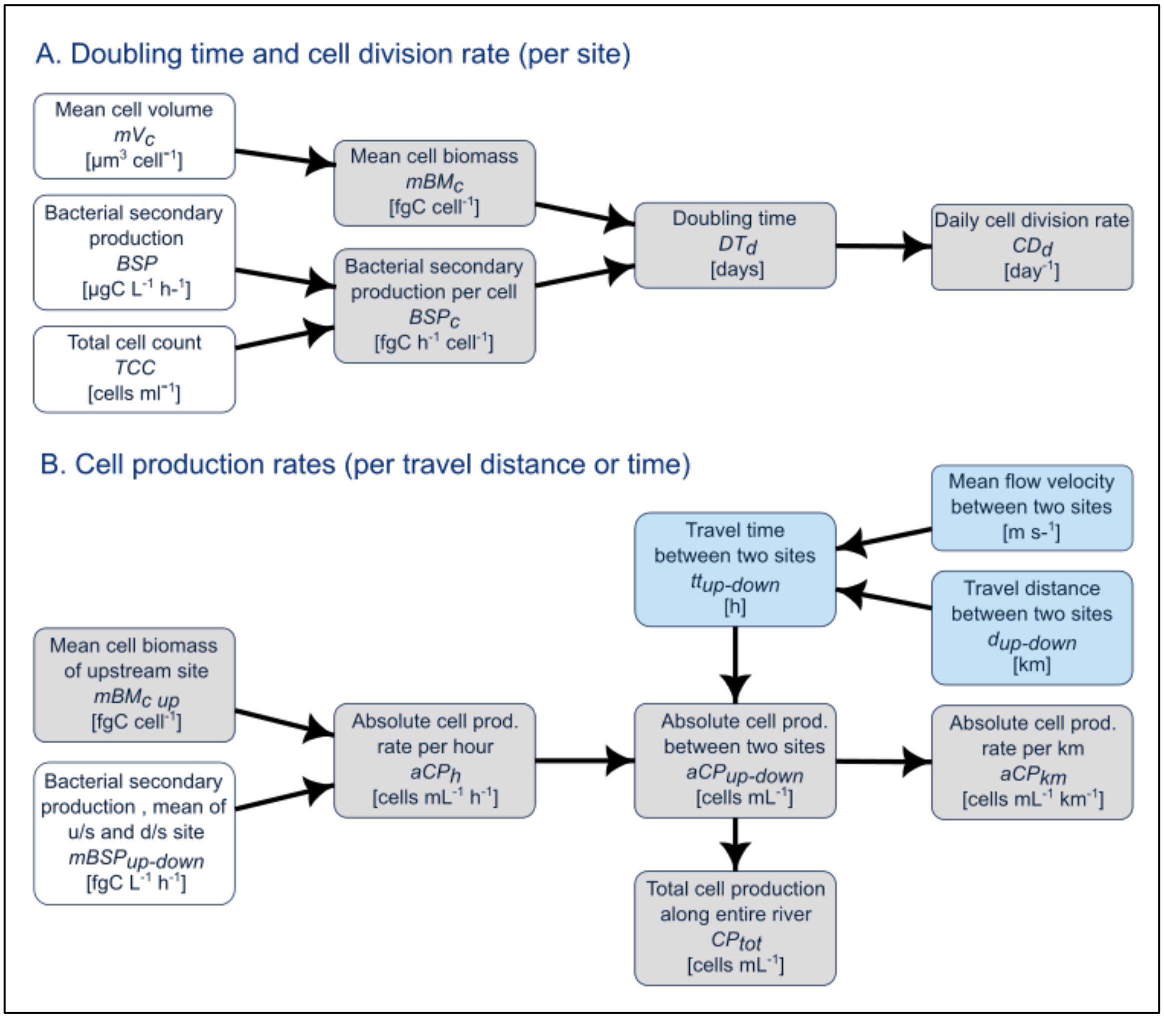


**Figure S2.** Schematic overview of calculation paths for bacterial cell division rates (**A**) and cell production rates (**B**). White background: measured microbiological parameters, grey background: calculated microbiological parameters, blue background: measured and calculated hydrological parameters. u/s: upstream, d/s: downstream.

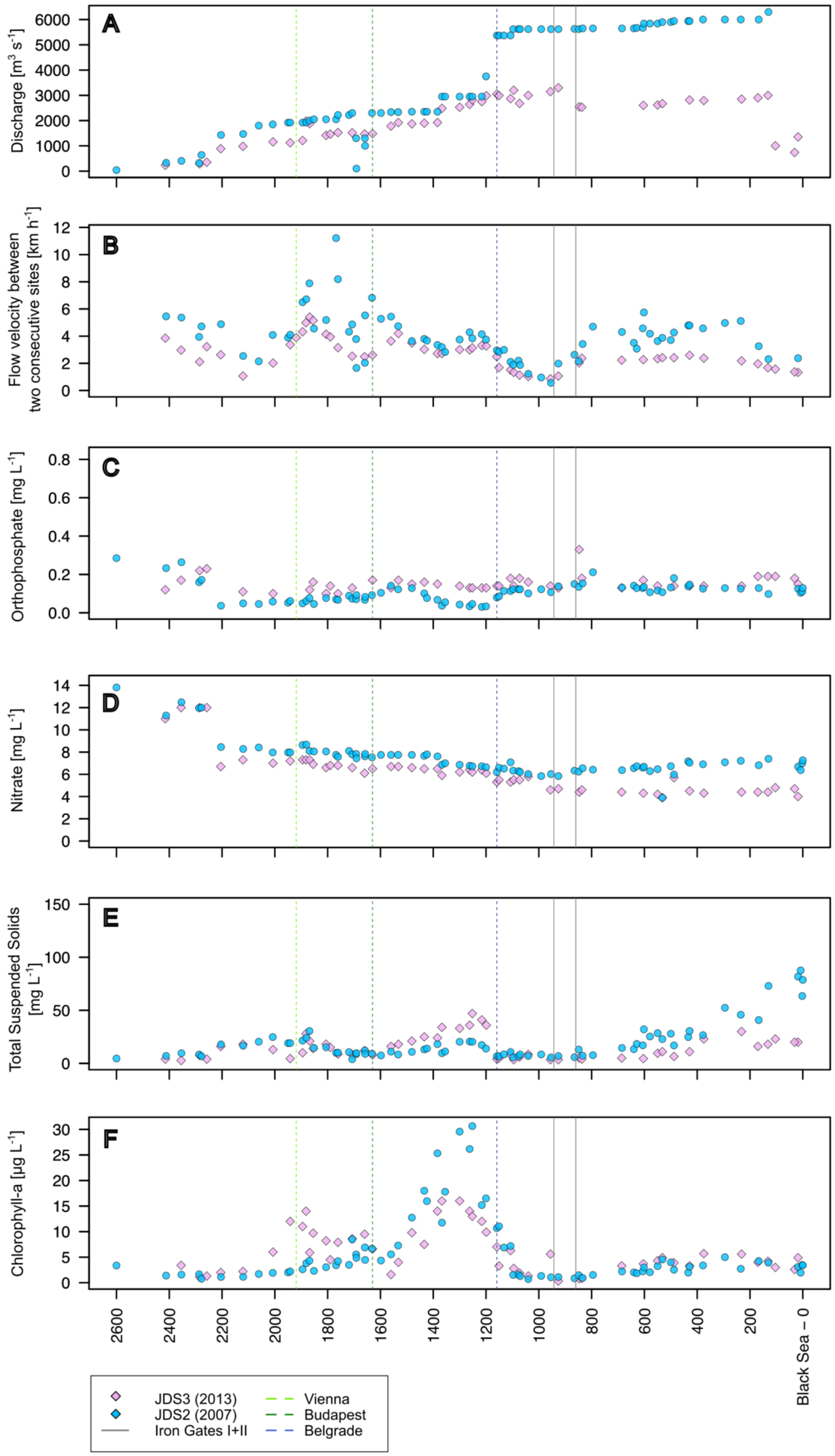
A
B
C
D
E
F
Discharge [m3 s-1]
Flow velocity between two consecutive sites [km h-1]
Orthophosphate [mg L-1]
Nitrate [mg L-1]
Total Suspended Solids [mg L-1]
Chlorophyll-a [µg L-1]
Black Sea – 0
JDS3 (2013)
JDS2 (2007)
Iron Gates I+II
Vienna
Budapest
Belgrade

**Figure S3.** Measured parameters along the longitudinal river transects for both Joint Danube Surveys (JDS) 2 & 3. n(JDS2) = 75, n(JDS3) = 54.

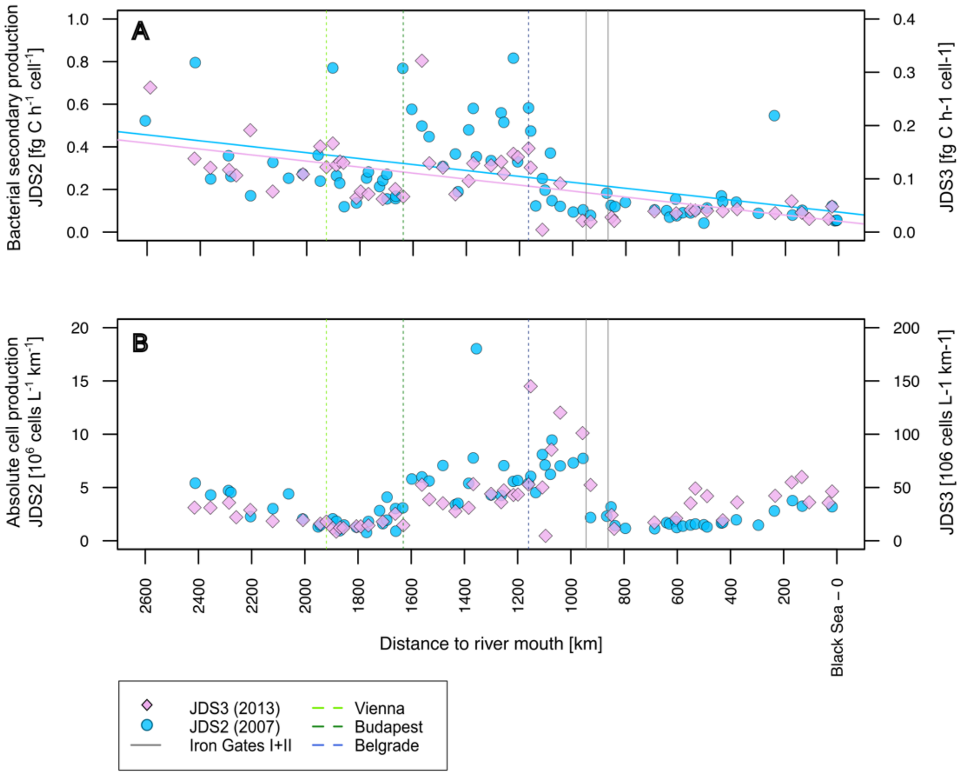


**Figure S4.** Trends in **(A)** cell-specific bacterial secondary production rates ($BSP_c$) and **(B)** absolute cell production rates per km between two sites ($aCP_{km}$) along the Danube River. n(JDS2) = 75, n(JDS3) = 54.

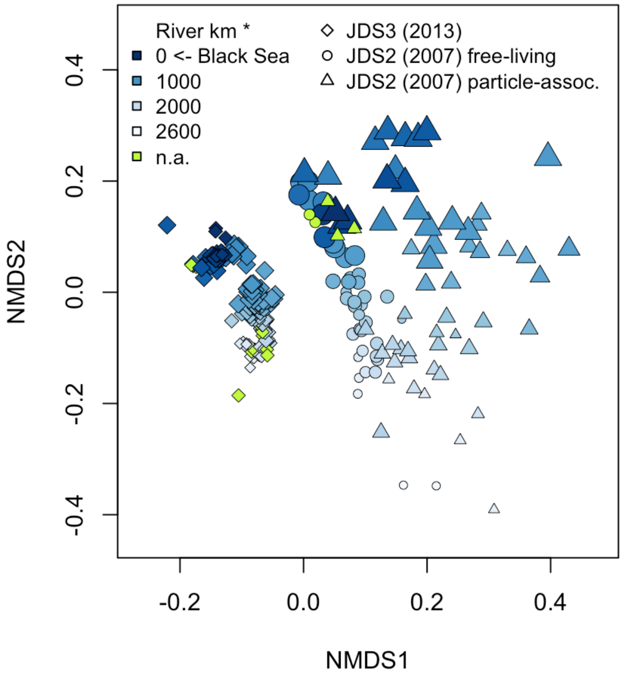


**Figure S5.** Non-metric multidimensional scaling (NMDS) of Bray-Curtis dissimilarities, illustrating the trends in longitudinal development of the bacterial communities along the river during the three studied datasets. The size of the symbols is proportional to the discharge at the respective site, the shade of blue indicates the location of the site along the river, from the most upstream site (white) towards the river mouth at the black see (rkm 0; blue).

**Table S1**: Summary statistics on covariation between environmental variables and the projections of bacterioplankton community samples in the Bray-Curtis based non-metric multidimensional scaling (NMDS) (**Fig. 3**). Values of covariation were calculated using the ‘envfit’ function included in the R-package ‘vegan’, fitting environmental vectors onto an ordination (Oksanen et al., 2013). Significance codes: *** ≤ 0.001, ** ≤ 0.01, * ≤ 0.05. † values reported in (Savio et al., 2015).

| **Parameter** | **Coefficient of determination ($R^2$)** | | |
|---|---|---|---|
| **Parameter** | **JDS3** | **JDS2 FL †** | **JDS2 PA †** |
| Distance to river mouth [river km] | 0.911*** | 0.840*** | 0.832*** |
| Catchment area [$km^2$] | 0.894*** | 0.766*** | 0.803*** |
| Cumulative dendritic stream lengths [km] | 0.885*** | 0.772*** | 0.807*** |
| Mean dendritic stream length [km] | 0.883*** | 0.799*** | 0.808*** |
| Median dendritic stream length [km] | 0.854*** | 0.752*** | 0.773*** |
| $NO_3^-$ | 0.709*** | 0.677*** | 0.538*** |
| Discharge (Q) [$m^3/s$] | 0.478*** | n. r. | n. r. |
| Chlorophyll a | 0.394*** | 0.016 | 0.396*** |
| Total suspended solids (TSS) | 0.258*** | 0.143* | 0.347*** |
| Mean flow velocity [m/s] | 0.236** | n. r. | n. r. |
| Max depth [m] | 0.229** | n. r. | n. r. |
| $PO_4^{3-}$ | 0.119 | n. r. | n. r. |
| Stream width [m] | 0.089 | ´n. r. | n. r. |

**Table S2.** Detailed information on ASVs showing noticeably high maximum growth or loss rates for the particle-asscociated size fraction of JDS2 (JDS2-PA), the free-living size fraction of JDS2 (JDS2-FL) and JDS3. The ASVs were selected based on a minimum ratio between maximum and minimum relative abundance of 100, a minimal maximum abundance of 1% in at least one dataset and a minimal prevalence in at least 20 samples.

| ASV | | | JDS2-PA | | JDS2-FL | | JDS3 | |
|---|---|---|---|---|---|---|---|---|
| | | | Relative abundance [%] | Absolute abundance [cells/L] | Relative abundance [%] | Absolute abundance [cells/L] | Relative abundance [%] | Absolu[illegible] abundan[illegible] [cells/L[illegible] |
| **Phylotypes with increasing abundance** | | | | | | | | |
| **ASV_3** | **hgcl_clade [Actinobacteriota]** | **min** | 0 | 0 | 0.3 | 8.1 | 1.9 | 98.8 |
| | | **max** | 4.4 | 139.6 | 5.2 | 186.2 | 16.2 | 2338.3 |
| **ASV_5** | **hgcl_clade [Actinobacteriota]** | **min** | 0 | 0 | 0.1 | 2.7 | 0.9 | 39.1 |
| | | **max** | 4.7 | 194.4 | 5.4 | 193 | 7.3 | 1567.4 |
| **ASV_15** | **hgcl_clade [Actinobacteriota]** | **min** | 0 | 0 | 0 | 0 | 0 | 0 |
| | | **max** | 2.6 | 82.7 | 2.1 | 54.5 | 6.9 | 1487.3 |
| **ASV_17** | **hgcl_clade [Actinobacteriota]** | **min** | 0 | 0 | 0 | 0 | 0 | 0 |
| | | **max** | 1.9 | 60.3 | 2 | 72 | 5.7 | 1167.1 |
| **ASV_21** | **CL500-29_marine_group [Actinobacteriota]** | **min** | 0 | 0 | 0 | 0 | 0 | 0 |
| | | **max** | 4 | 150.4 | 4.8 | 106.8 | 9.1 | 1714.2 |
| **Phylotypes with decreasing abundance** | | | | | | | | |
| **ASV_25** | **Limnohabitans [Proteobacteria]** | **min** | 0 | 0 | 0 | 0 | 0 | 0 |
| | | **max** | 1.3 | 30.3 | 1.2 | 35.3 | 3.1 | 223.7 |
| **ASV_27** | **hgcl_clade [Actinobacteriota]** | **min** | 0 | 0 | 0 | 0 | 0 | 0 |
| | | **max** | 2.3 | 53.5 | 1.2 | 29.8 | 4.4 | 299.5 |
| **ASV_37** | **Sphingorhabdus [Proteobacteria]** | **min** | 0 | 0 | 0 | 0 | 0 | 0 |
| | | **max** | 1.5 | 46.9 | 1.7 | 57.7 | 2 | 186.3 |
| **ASV_55** | **Sediminibacterium [Bacteroidota]** | **min** | 0 | 0 | 0 | 0 | 0 | 0 |
| | | **max** | 0.9 | 20.9 | 1 | 24.3 | 1.3 | 160 |
| **ASV_71** | **Limnohabitans [Proteobacteria]** | **min** | 0 | 0 | 0 | 0 | 0 | 0 |
| | | **max** | 0.5 | 10.2 | 0.6 | 10.9 | 1.1 | 84.4 |
| **ASV_73** | **CL500-29_marine_group [Actinobacteriota]** | **min** | 0 | 0 | 0 | 0 | 0 | 0 |
| | | **max** | 0.8 | 23.2 | 0.7 | 14.6 | 1.9 | 154.3 |
